# Co-expression of calcium channels and delayed rectifier potassium channels protects the heart from proarrhythmic events

**DOI:** 10.1101/659821

**Authors:** Sara Ballouz, Melissa M Mangala, Matthew D Perry, Stewart Heitmann, Jesse A Gillis, Adam P Hill, Jamie I Vandenberg

**Affiliations:** Stanley Institute for Cognitive Genomics, Cold Spring Harbor Laboratory, Woodbury, NY 11797, USA; Victor Chang Cardiac Research Institute, Darlinghurst, NSW 2010, Australia; University of NSW, Sydney. Kensington, NSW 2052, Australia

**Keywords:** Ion channels, calcium channels, potassium channels, co-expression, gene modules, meta-analysis, cardiac action potential, early afterdepolarization, arrhythmia, drug-induced long QT syndrome

## Abstract

Cardiac electrical activity is controlled by the carefully orchestrated activity of more than a dozen different ion conductances. Yet, there is considerable variability in cardiac ion channel expression levels both within and between subjects. In this study we tested the hypothesis that variations in ion channel expression between individuals are not random but rather there are modules of co-expressed genes and that these modules make electrical signaling in the heart more robust.

Meta-analysis of 3653 public RNA-Seq datasets identified a strong correlation between expression of *CACNA1C* (L-type calcium current, *I*_CaL_) and *KCNH2* (rapid delayed rectifier K^+^ current, *I*_Kr_), which was verified in mRNA extracted from human induced pluripotent stem cell-derived cardiomyocytes (hiPSC-CM). *In silico* modeling, validated with functional measurements in hiPSC-CM, indicates that the co-expression of *CACNA1C* and *KCNH2* limits the variability in action potential duration and reduces susceptibility to early afterdepolarizations, a surrogate marker for pro-arrhythmia.

**Impact Statement:** Coexpressed levels of potassium and calcium ion channel genes in the heart encode more robust cardiac electrophysiology and provide insights into genetic basis of arrhythmic risk

## Introduction

Robust electrical signaling in the heart is critical for co-ordinating the efficient pumping of blood around the body. Failure of cardiac electrical signaling, even for just a few minutes, can have fatal consequences, with sudden cardiac death accounting for up to 10% of deaths in our community (1). Despite decades of research, predicting in advance who is more or less susceptible to sudden cardiac death remains challenging (2).

The action potential (AP) of excitable cells, such as cardiac myocytes and neurons, reflects the orchestrated activity of at least a dozen distinct ion channels and electrogenic transporters (3,4). In such complex systems, both theoretical and experimental studies have shown that there is considerable inter-individual variability in the combinations of molecular input parameters that can produce very similar integrated outputs (5,6). This has led to a paradigm shift in computational cardiology that relies not on generating idealized outputs based on mean data but rather development of populations of models that account for the observed variability in molecular inputs (7,8). Such models are already proving useful for interrogating inter-individual variability in response to pathological stimuli (9–11). Our challenge now is to discern the underlying essence of these complex systems (12,13) so that we may then make rational interventions to treat pathology that takes into account inter-individual variation. Specifically, are there underlying principles regulating cardiac electrical activity that can provide insights into why some people are more susceptible to sudden cardiac death in response to pathological stimuli, such as drug block of the rapid delayed rectifier potassium channel (*I*_Kr_) which is the underlying basis of drug-induced long QT syndrome (di-LQTS) (14).

A common approach to discern patterns in multi-dimensional biological problems has been to look for co-expression networks (13,15). Co-expression networks are known to encode functional information (16), with co-expression reflecting co-regulation and co-functionality (17,18). To help identify robust co-expression modules, it is helpful to use meta-analytical approaches, which aggregate large numbers of individual networks across multiple independent experiments to average away noise and reinforces those correlations that reflect real signals (19–21).

Here, we have used meta-analytic co-expression analysis in large scale human gene expression data sets to identify modules of co-expressed ion channel genes which were then used to constrain population models of cardiac electrical activity. These models were then used to test the hypothesis that co-expression of repolarization and depolarization currents helps prevent irregular APs in human cardiac myocytes that are exposed to pro-arrhythmic stimuli. We show that tight coupling of current densities for the L-type calcium current (*I*_CaL_) and the rapid component of the delayed rectifier potassium current (*I*_Kr_) reduced the emergence of pro-arrhythmic early afterdepolarizations (EADs) and this protection persisted in the face of highly variable expression of other ion channels, as well as in the presence of pharmacological block of *I*_Kr_ which is a potent pro-arrhythmic stimulus (22). A very important prediction to arise out of our modelling studies is that in the context of drug block of *I*_Kr_, those patients with low expression of *I*_CaL_ and *I*_Kr_ will exhibit the greatest prolongation of repolarization duration but patients with high expression of *I*_CaL_ and *I*_Kr_ will experience more EADs and are therefore more likely to be susceptible to ventricular arrhythmias.

## Results

The shape and duration of action potentials in cardiac myocytes are determined by the orchestrated activity of voltage-gated sodium, calcium and potassium channels, as well as a series of electrogenic transporters that regulate intracellular ion concentrations. These channels, transporters and related intracellular calcium handling proteins are encoded by a few dozen genes, sometimes referred to as the rhythmonome (23) (a full list is provided in Figure 1 source data 1).

To determine whether there were any co-expression patterns among the rhythmonome genes we first undertook an untargeted screen for possible expression correlation patterns in publicly available RNA-seq data sets (**Figure 1**). Ranked correlation coefficients from an aggregate co-expression network that contain data from 3653 samples are illustrated in the heatmap in **Figure 1A**. High ranked correlations indicate similarity of transcriptional profiles between the genes. A clustering analysis, as shown by the dendrogram in **Figure 1A**, groups genes according to their correlation similarities, as defined by the Spearman’s correlation coefficients (see color code in **Figure 1A**). There is a large cluster of yellow/green squares in the bottom left corner indicating that there are significant levels of correlation amongst many of the genes. Furthermore, within this large cluster there are two sub-clusters. The cluster of yellow-green squares corresponding to 13 genes in the bottom left corner (red dashed box in **Figure 1A**) encode for proteins that regulate calcium fluxes as well as the transient outward K^+^ current (*KCND3*), which helps to maintain the plateau potential at a level that maximizes calcium influx (24). A second sub-cluster, in the upper right quadrant of the main cluster, encompasses 10 genes (black dashed box in **Figure 1A)**, including *KCNH2*, *SCN5A*, *KCNJ12* and *KCNIP2*, that encode for ion channel proteins important for regulating action potential duration (APD).

**Figure 1:**
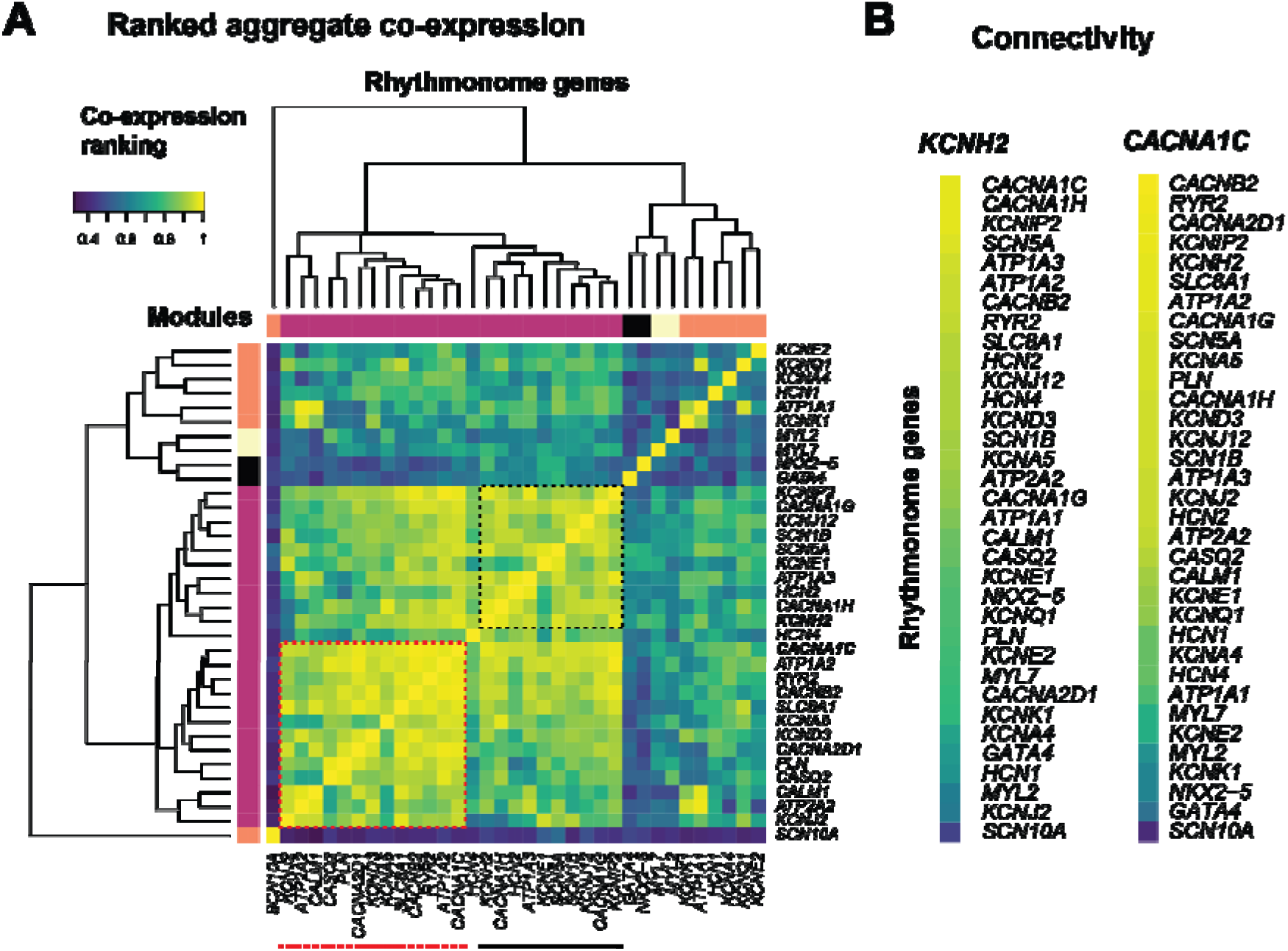
Cardiac ion channel and calcium handling protein co-expression connectivity analysis from published RNA-seq datasets. The full list of RNA-seq experiments interrogated are summarized in Figure 1 source data 1). **A.** Sub-network of co-expression ranking across 35 genes encoding for cardiac ion channels or calcium homeostasis proteins. Blue: low ranked correlations, yellow: high ranked correlations. The dendrogram illustrates the modules of genes with high levels of similarity in their transcriptional profiles. Red and black dashed boxes highlight subnetworks of correlated genes **B.** Connectivity of genes to **KCNH2** (encodes for *I*_Kr_) and **CACNA1C** (encodes for *I*_CaL_).

Overall, the connectivity within the set of rhythmonome genes is fairly high in comparison to their connectivity to all other genes in the co-expression network (node degree analysis, p~5.7e−14), with a central gene being *CACNA1C*, the gene encoding the alpha subunit of the L-type calcium channel (see **Figure 1, figure supplement 1**). Although *CACNA1C* clusters in the group of calcium handling genes in the bottom left quadrant of the main cluster in **Figure 1A**, it also shows high correlation with the cluster of ion channel genes in the top right quadrant of **Figure 1A**. In a similar fashion, the *KCNH2* gene, which encodes for *I*_Kr_, is included in the ion channel cluster but also shows moderate-high correlations with a portion of the calcium handling genes in the bottom left quadrant. The highest ranked co-expression partners for *CACNA1C* and *KCNH2* are highlighted in **Figure 1B**.

The vast majority of the public RNA-Seq datasets included in our analyses were not heart specific. It is, however, noteworthy that there are no strong correlations between the expression of any of the individual ion channel or calcium handling genes and the cardiac-specific markers included in our analyses: *GATA-4*, *NKX2-5*, *MYL2* and *MYL7*. For example, the cardiac marker genes are towards the bottom of the lists in **Figure 1B**. This suggests that the correlations within the set of rhythmonome genes are not simply a reflection of cardiac specific expression but rather represent intrinsic correlations.

To investigate whether the meta-analytic co-expression patterns observed in **Figure 1** were also seen in human heart cells, we extracted mRNA from human induced pluripotent stem cell-derived cardiomyocytes (hiPSC-CMs) (25) obtained from 10 patients with no known heart disease. All samples contained high levels of cardiac marker genes (*MYL7*, *GATA4* and *NKX2*.5) **(Figure 2A)**. The levels of expression of most rhythmonome genes showed variations between the samples that spanned approximately an order of magnitude. However, similar to the generic tissue RNA-Seq datasets, there were modules of co-expressed genes (e.g., see dashed box in bottom left quadrant of **Figure 2B**). Most of the 13 genes in the cluster in **Figure 2B** are present in the modules highlighted by the black and red dashed boxes in **Figure 1A**. Conversely, many of the ion channel genes contained in the modules highlighted in **Figure 1** are not present in the module in **Figure 2** (e.g. *KCNJ2*/*KCNJ12*, which encodes for *I*_K1_; *KCND3*, which encodes for *I*_To_; and *SCN5A*, which encodes for *I*_Na_). These genes are all known to be expressed at lower levels in embryonic hearts and so it is unsurprising that they are not well expressed in the hiPSC-CM lines (26).

**Figure 2:**
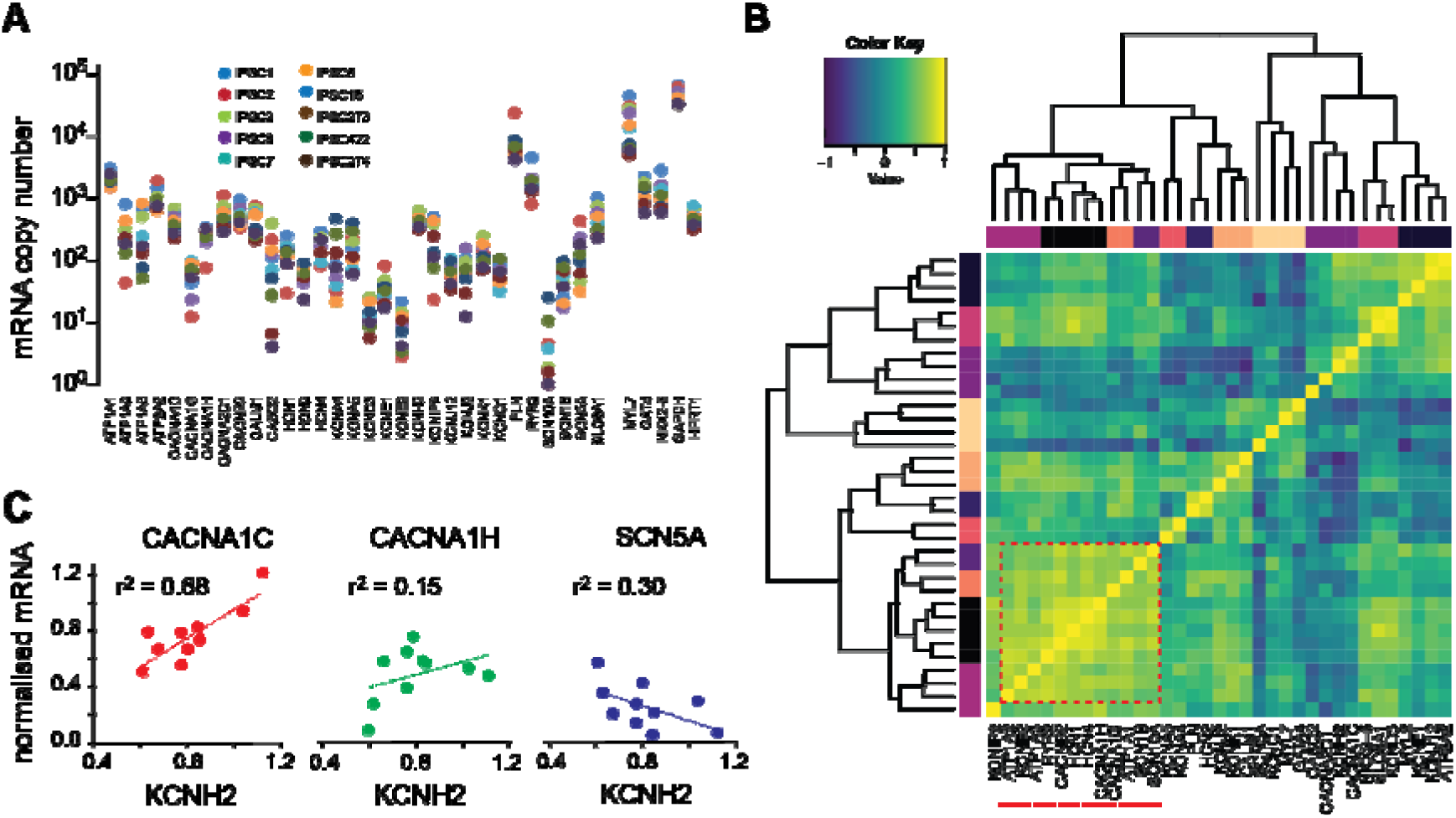
HiPSC-CM mRNA correlation analysis. **A.** plot of all mRNAs (log axis) for 10 hiPSC-CM lines. Note that there are high levels of expression of cardiac markers (*MYL7* as well as *NKX2-5* and *GATA4*) in all cell lines. Also, there is less variability in the levels for the housekeeping genes (*GAPDH* and *HPRT1*) compared to that seen for the ion channels and calcium handling proteins. The mRNA data for panel A are provided in Figure 2 -Source Data 1 (GEO: TBA). **B.** Correlation matrix for hiPSC-CM mRNA. The red dashed box highlights the module with the highest correlations. **C.** Correlation plots for normalised mRNA expression *KCNH2* versus genes that encode the major depolarizing currents (*CACNA1C*, *CACNA1H* and *SCN5A*). To account for any differences in mRNA loading between runs, levels of expression have been normalized to GAPDH.

We next looked to see if there were any specific relationships between genes encoding depolarization and repolarization currents, within the hiPSC-CM expression profiles. In **Figure 2C**, we have plotted the expression of *KCNH2* versus the genes that encode for the depolarization currents that showed the highest levels of correlation with expression of *KCNH2* in the generic tissue datasets (see **Figure 1B**, i.e., *CACNA1C*, *CACNA1H* and *SCN5A*). The most notable correlation that was observed in the hiPSC-CMs was that between *KCNH2* and *CACNA1C*; r^2^ = 0.68 (**Figure 2C**).

As the most robust relationship that we observed in both the generic tissue sets and the hiPSC-CM lines was the co-expression of *KCNH2* and *CACNA1C*, we focused on this pair for our subsequent studies. To investigate whether co-expression modules of ion channel genes might influence integrated cardiac electrical function, we used an *in silico* approach (**Figure 3**). We used the O-Hara-Rudy model of the human ventricular myocyte (27) with the baseline conductance scalars optimized as previously described (28). First, we simulated a population of 1000 human cardiac APs where the ion conductances for each cell were scaled randomly (see **Figure 3A. Figure 3 movie supplement 1**).

**Figure 3:**
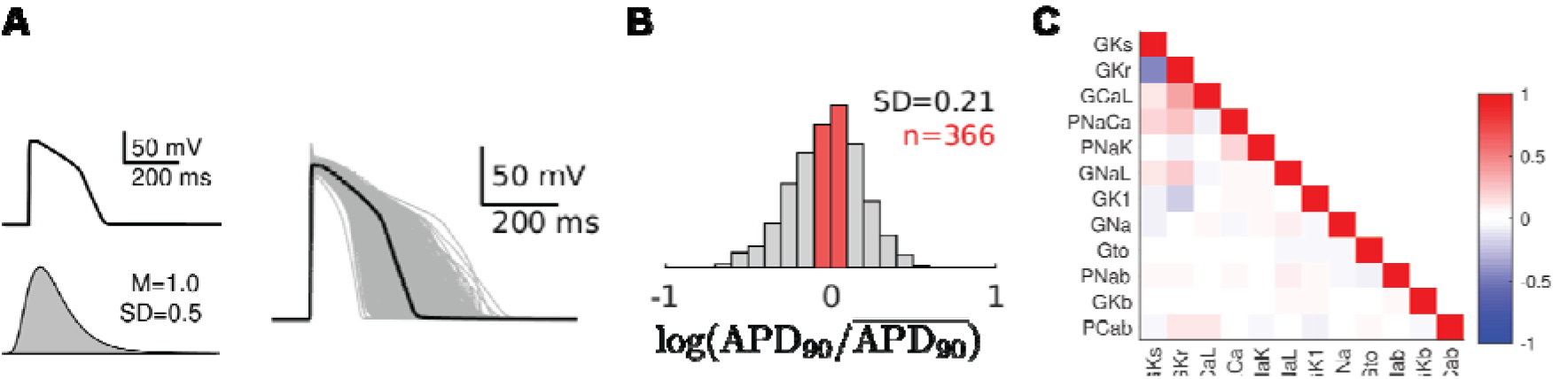
*in silico* predictions of conductance scalars that give APD_90_ values in the normal range. **A.** Baseline AP model and frequency histogram of scalars (log normal distribution with a mean of 1 and SD of 0.5) used to generate the population of 1000 APs with the baseline AP (all scalars set to 1.0) shown in black. The last 4 beats for each of the population of 1000 APs are shown in the **Figure 3 movie supplement 1** (see below). **B.** Frequency histogram of APD_90_ values for the 1000 simulations with those falling within ±20% of the mean value shown in red. **C.** correlation matrix for sets of conductance scalars that gave APD_90_ values within ±20% of the mean value

The baseline action potential produced by this model (black trace in right hand panel of **Figure 3A**) has a duration at the point of 90% repolarization (APD_90_) of 264 ms. Whereas, the APD_90_ values in the cells with randomized conductance scalars ranged from ~120 ms to ~500 ms (**Figure 3A**, right panel). We next selected those cells with APD_90_ values that fell within 20% of the mean value (red bars, **Figure 3B**) to determine if there were any patterns of ion channel co-expression that could contribute to keeping the APD_90_ within this narrow-selected range. The correlation matrix of the conductance scaling factors for the selected cells (**Figure 3C**) reveals a positive correlation between *G*_Kr_ and *G*_CaL_ (R=0.36) as well as an inverse correlation for *G*_Kr_ and *G*_Ks_ (R=−0.46). The positive correlation between the conductance scalars *G*_Kr_ and *G*_CaL_ in the *in silico* modelling dataset (**Figure 3**) suggests that the correlation seen between *CACNA1C* and *KCNH2* mRNA expression in both the public RNA-Seq datasets (**Figure 1**) and the hiPSC-CM dataset (**Figure 2**) would contribute to reducing the population variability in APD_90_ values.

We next repeated our previous simulation but forced the conductance scaling factors for *I*_Kr_ and *I*_CaL_ to be identical in each cell (denoted co-expression in **Figure 4**). The scaling factors for all other conductances were allowed to vary independently. The distribution of APD_90_ values for both independent and co-expression cell populations becomes broader as the level of variability is increased (**Figure 4A-C**). However, the spread of APD_90_ values in the cells with identical *G*_CaL_-*G*_Kr_ scalars is always narrower than in the cell populations with independent *G*_CaL_-*G*_Kr_ scalar values. For example, in the case of **Figure 4B**, the variance of the APD_90_ values was 0.026 for the co-expression dataset but 0.046 for the independent dataset (see **Figure 4 figure supplement 1**). Another notable feature of the data in **Figure 4C** is that early afterdepolarizations (EADs) begin to appear in the cell population with independent scalars when the scalar variability, σ^2^, exceeds 0.20 (also see **Figure 4 Figure supplement 1**). The number of cells with an EAD are indicated in parentheses above each distribution in **Figure 4C**.

**Figure 4:**
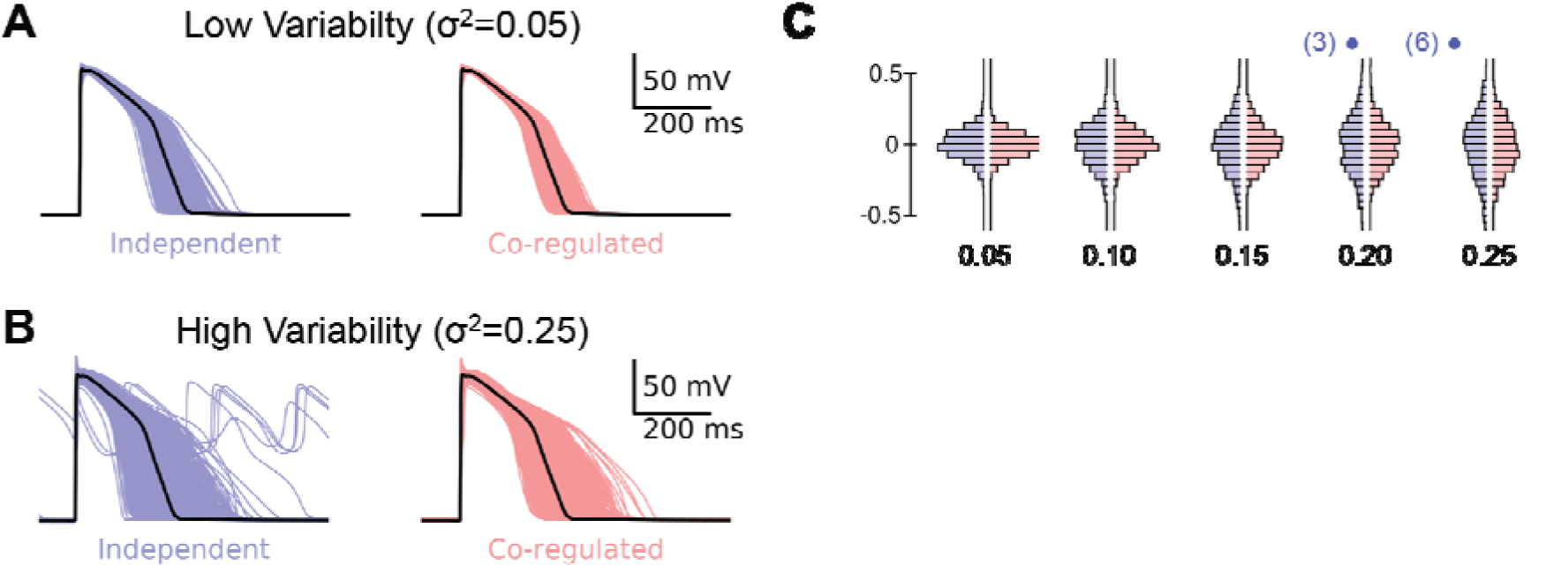
Impact of *I*_Kr_-*I*_CaL_ co-expression and conductance scalar variability on APD_90_ variability. APs for independent (blue) and co-expression of *G*_Kr_-*G*_CaL_ (red) for **A.** low scalar variability (σ^2^=0.05) and **B.** high scalar variability (σ^2^=0.25). Note the presence of EADs in 6 of the APs in the independent group with σ^2^ = 0.25. The last 4 beats for each of the population of 1000 APs are shown in the **Figure 4 movie supplement 1**. **C.** Histograms of log(APD_90_/mean APD_90_) distributions for co-expressed (red) or independent (blue) *G*_Kr_-*G*_CaL_ simulations with σ^2^ = 0.05, 0.1, 0.15, 0.2, 0.25. The numbers in parentheses above the 0.2 and 0.25 groups indicate the number of EADs in each independent scalars group. APs and incidence of EADs for an extended range of variances are shown in **Figure 4 figure supplement 1**.

We next investigated whether coupling of the conductance scalars for *G*_Kr_ and *G*_CaL_ influenced the generation of EADs in response to a pathological stimulus. Specifically, we investigated how cells respond to drug block of *I*_Kr_ which is the underlying cause of di-LQTS (22). Populations of action potentials obtained for independent and co-expression populations with *I*_Kr_ block of 0%, 50% and 80% are illustrated in **Figure 5**. As the extent of *I*_Kr_ block is increased (from A to C), the proportion of action potentials producing EADs increases. It is also clear that at lower levels of *I*_Kr_ block, EADs were more frequent when *G*_CaL_ and *G*_Kr_ scalars were modulated independently (see **Figure 5B** and inset to **Figure 5D**). However, the proportion of simulations developing EADs in both the independent and co-expression populations becomes similar when the extent of *I*_Kr_ block exceeds 80% (**Figure 5D**).

**Figure 5:**
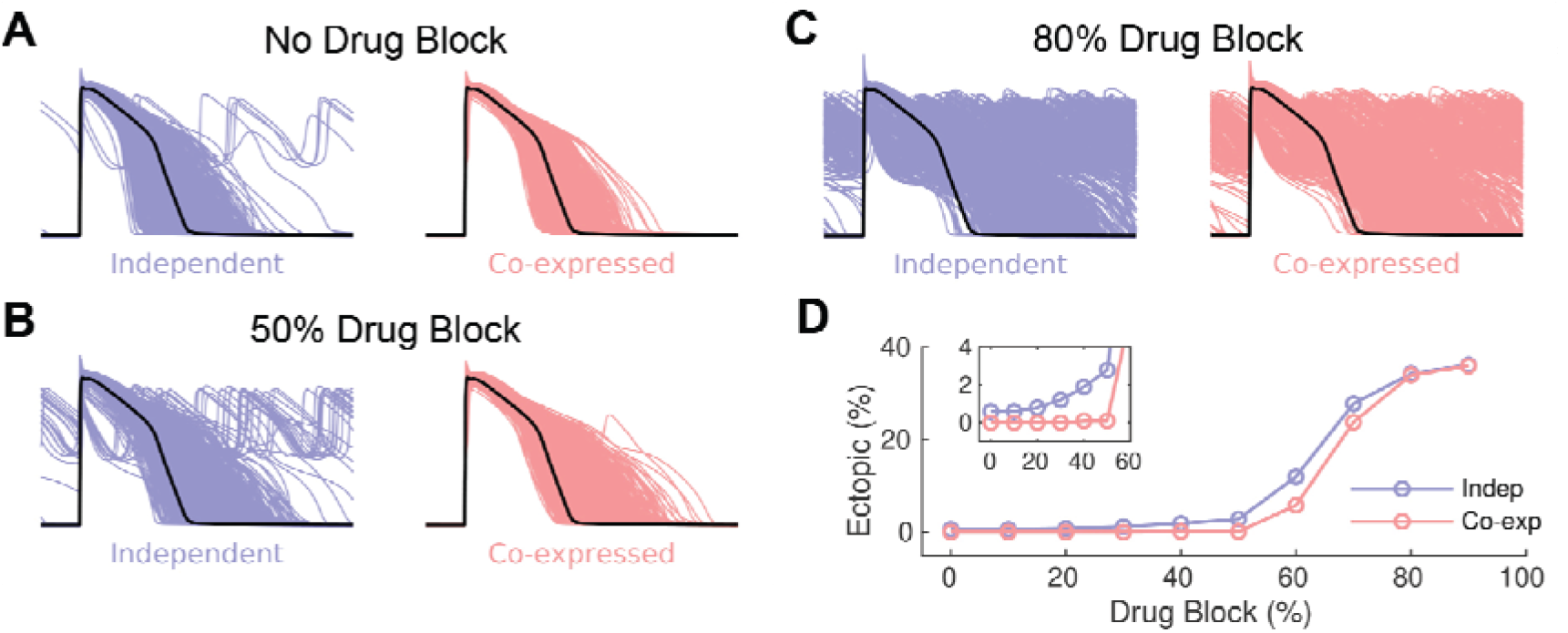
Impact of *G*_Kr_-*G*_CaL_ co-expression on response to drug block of *I*_Kr_. APs for co-expressed *G*_Kr_-*G*_CaL_ (red) and independent (blue) *G*_Kr_-*G*_CaL_ populations, with scalar variance σ^2^ = 0.25, for **A.** control, **B.** 50% *I*_Kr_ block and **C.** 80% *I*_Kr_ block. **D.** plot of % of AP simulations with EADs versus % *I*_Kr_ block (co-expression, red and independent, blue). The inset in panel D shows an expanded view of the % EADs at small levels of *I*_Kr_ block.

It is well established that the extent of repolarization prolongation seen in patients prescribed drugs that block *I*_Kr_ is highly variable and only a subset of patients exposed to drugs that block *I*_Kr_ (29), or with a mutation causing 50% loss of *I*_Kr_ function (30), will develop life threatening arrhythmias. This is consistent with the prediction made by our simulated drug block experiments shown in **Figure 5**. We therefore asked whether the data from the co-expression datasets could tell us anything about what factors might predispose to lesser or greater extent of AP prolongation and/or the development of EADs in the presence of a drug that blocks *I*_Kr_. Analysis of the subset of scalars within the co-expression dataset that produced the 50 longest APs without EADs, compared to the subset of scalars that resulted in APs with EADs, is illustrated in **Figure 6A and 6B** respectively. Notably, the longest APs without EADs had low *G*_CaL_ scalars (and hence low *G*_Kr_ scalars before addition of drug block). Conversely, the APs that developed EADs had higher *G*_CaL_ and *G*_Kr_ scalars. In **Figure 6C**, we have plotted the APD_90_ values for cells in the highest (red) and lowest (blue) quartiles of *G*_CaL_ - *G*_Kr_ scalars. As expected, the low *G*_CaL_ group showed longer APD_90_ values on average compared to the high *G*_CaL_ group (see the continuous lines in **Figure 6C**). Furthermore, for the 70% *I*_Kr_ block scenario, 44% of the high *G*_CaL_ group have developed EADs whereas only 7% of the low *G*_CaL_ group have developed EADs (compare red and blue bars at 70% drug block in **Figure 6C**). Thus, higher *G*_CaL_ is associated with a greater risk of developing EADs in response to moderate levels of *I*_Kr_ block. A similar pattern of results was observed when *G*_CaL_ and *G*_Kr_ were allowed to vary independently, except that in this scenario the difference between the high *G*_CaL_ and low *G*_CaL_ groups was even more dramatic at lower levels of *I*_Kr_ block (see **Figure 6, figure supplement 1**).

**Figure 6:**
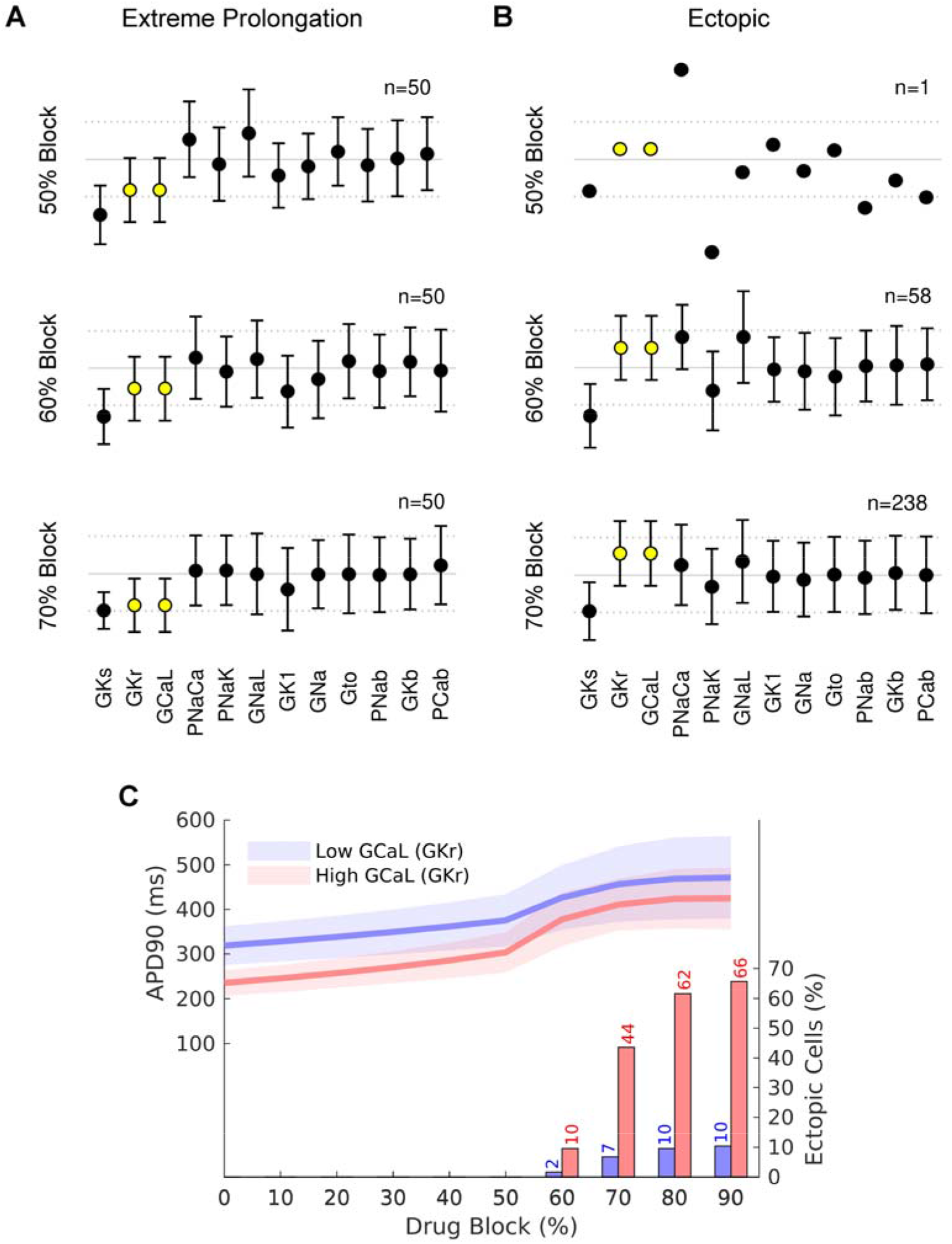
High *G*_CaL_ increases risk of EAD formation at moderate *I*_Kr_ block. Scalars (mean ± SD) for the population of cells with *G*_Kr_-*G*_CaL_ co-expression. **A.** 50 cells with the longest APD_90_ values and no EAD with increasing % *I*_Kr_ block. **B.** Scalars (box and whisker plots) for cells with EADs with increasing % *I*_Kr_ block. The *G*_CaL_ and *G*_Kr_ scalars are highlighted in yellow. **C.** APD_90_ values (left axis) for the highest (red) and lowest (blue) quartile of cells according to baseline *G*_CaL_ scalar. The continuous lines show the mean value and shaded area shows ±1 SD). The percentage of EADs in each quartile is shown as columns (see axis on right side of graph). The corresponding plots for the population of cells with independent *G*_Kr_-*G*_CaL_ scalars are show in the **Figure 6 figure supplement 1.**

A corollary of our prediction that patients with high *G*_CaL_ are more susceptible to EADs when exposed to a drug that blocks *I*_Kr_, is that co-administration of a drug that blocks *I*_CaL_ would reduce the incidence of EADs. Magnesium, which is used in the acute management of di-LQTS (14), is a weak calcium channel blocker. Raising plasma [Mg^2+^] from 1.5 to 2.5 mM would be expected to inhibit *I*_CaL_ by ~20% (31), conversely reducing [Mg^2+^] from 1.5 to 0.5 mM would be expected to increase *I*_CaL_ by ~20%. When we increased *I*_CaL_ by 20%, the incidence of EADs in the high *G*_CaL_ group increased from 10% to 16% for the 60% *I*_Kr_ block simulation and from 44% to 58% for the 70% *I*_Kr_ block simulations (see **Figure 7**). A drug that inhibited *I*_CaL_ by 20% caused a modest decrease in the percentage of cells with EADs and reduction of *I*_CaL_ by 50% had a correspondingly larger effect, for example, reducing EADs from 44% to 12% in the 70% *I*_Kr_ block scenario (**Figure 7**).

**Figure 7:**
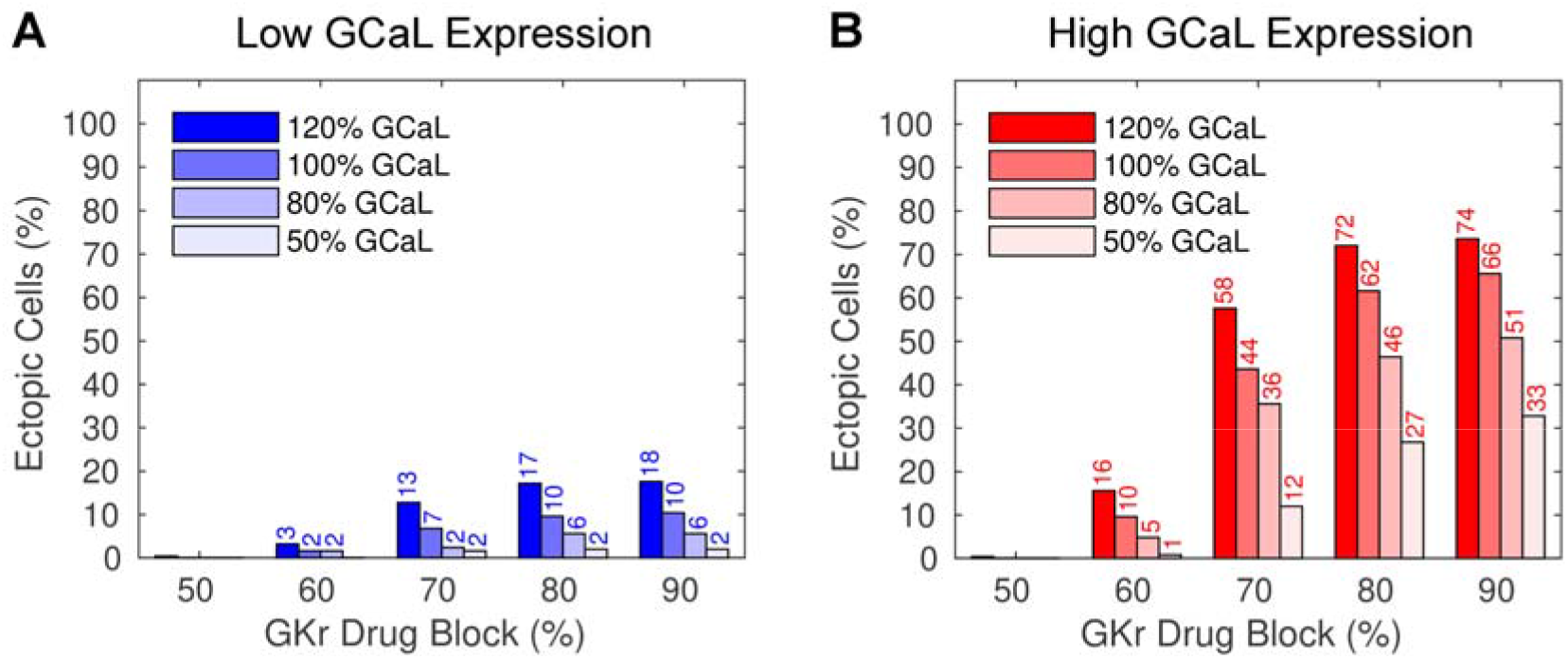
Effect of modifying *I*_CaL_ on incidence of EADs in response to *I*_Kr_ drug block. *I*_CaL_ was increased 20% to mimic hypomagnesaemia, reduced 20% to mimic hypermagnesaemia (see text for details), or reduced 50% to mimic administration of a calcium channel blocker. In both the quartile of cells with the lowest *G*_CaL_ scalars (blue, **A**) and the quartile of cells with the highest *G*_CaL_ scalars (red, **B**) the incidence of EADs increased when *G*_CaL_ was enhanced and decreased when *G*_CaL_ was decreased. The observed differences were most pronounced with moderate (60-70%) *I*_Kr_ drug block.

To determine whether our *in silico* predictions could be validated in cardiac myocytes we investigated whether the extent of *I*_Kr_-block induced prolongation of repolarization seen in cardiac myocytes derived from different hiPSC lines were correlated with the levels of *KCNH2* gene expression. We measured the duration of repolarization from extracellular field potentials (FPD) recorded from monolayers of hiPSC-derived cardiac myocytes paced at 1 Hz (see **Figure 8**). Cisapride (200 nM), a highly selective *I*_Kr_ blocker (32), caused a prolongation of repolarization that varied between 17.9 ± 7.4 ms (mean ± SD, n=12) for the least sensitive cell line to 98 ± 41 ms (mean ± SD, n=11) for the most sensitive cell line (**Figure 8A**). There was a significant inverse correlation between the extent of FPD prolongation and the [*KCNH2*-mRNA] measured for each cell line (**Figure 8B**). Furthermore, the extent of FPD prolongation caused by Cisapride in each cell line was almost exactly reversed by Nifedipine (50 nM), a highly selective *I*_CaL_ blocker (32) (Figure 8A).

**Figure 8:**
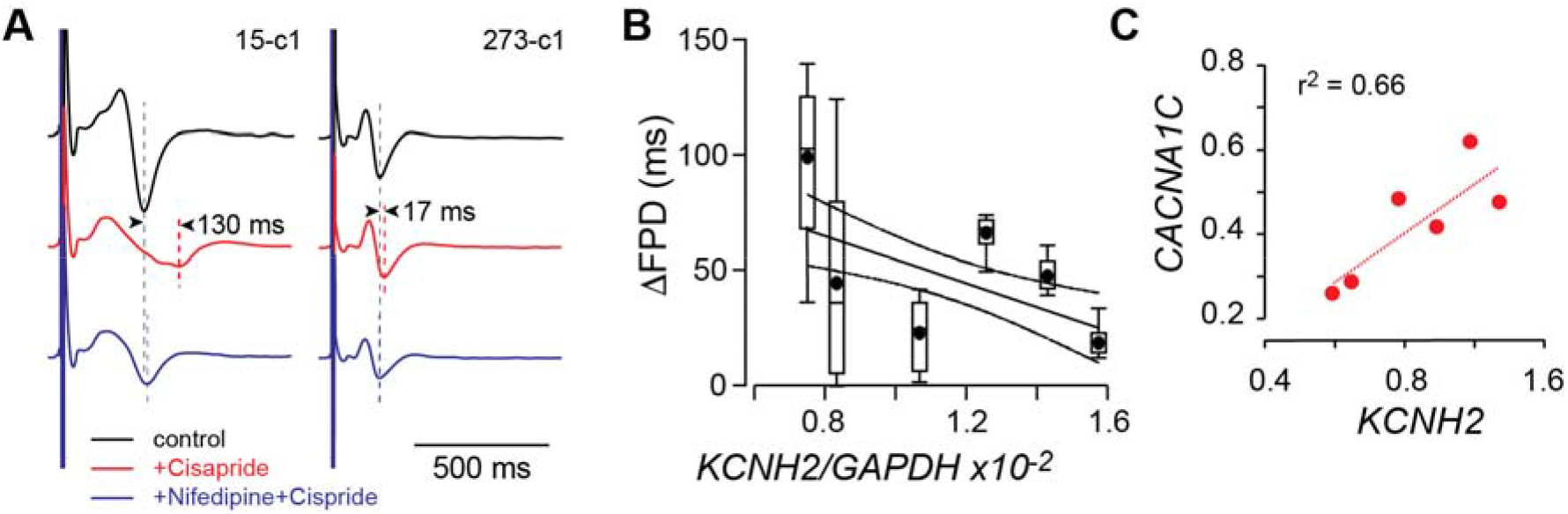
Effect of *I*_Kr_ and *I*_CaL_ inhibition on repolarization duration in iPSC-CM. **A.** Example extracellular electrograms recorded from two separate hiPSC-CM lines under control (black) conditions, after 5 minutes perfusion with cisparide (200 nM, red)followed by 5 minutes perfusion with cisapride (200 nM) + nifedipine (50 nM, blue). In cell line 15-c1 there was a 130 ms prolongation in field potential duration (FPD) which almost completely reversed on addition of nifedipine. In cell line 273-c1 there was a much smaller prolongation of 17 ms with addition of Cisapride that again reversed on the addition of Nifedipine. **B.** Summary of ΔFPD for 6 cell lines plotted against the level of *KCNH2* mRNA (normalised to GAPDH expression). Data shown as box and whiskers (median, box: 25%-75%, extremities: 5%-95% with the mean shown as a filled circle, n=7-12 replicates for each cell line). The straight line of best fit is shown as a solid line with 95% confidence intervals shown as dashed lines; the slope of the line is significantly different to zero (F-test, p=0.0010). The micro-electrode array data for panels A&B are provided in Figure 8 - Source Data 1. **C.** Plot of normalised mRNA expression for *KCNH2* versus *CACNA1C* for the cell lines used in the experiments summarized in panel B. To account for any differences in mRNA loading between runs, levels of expression have been normalized to GAPDH. The mRNA data for panels B&C are provided in Figure 8 -Source Data 2 (GEO: TBA).

## Discussion

Cardiac electrical activity is regulated by the interdependent activity of a plethora of ion channels, transporters and calcium handling proteins (4). Understanding the precise details of how these conductances interact to control the rhythm of the heart has been an enduring source of fascination. In this study, we have used meta-analytical techniques to interrogate the large numbers of RNASeq datasets that have been deposited in public access databases, to look for patterns of co-expressed genes that might help decipher the control of cardiac electrical activity. The most important pair of co-expressed genes, identified both in the public RNA Seq datasets (**Figure 1**) and in heart cells derived from hiPSC lines (**Figure 2**), was that of *CACNA1C* (*I*_CaL_) and *KCNH2* (*I*_Kr_). Using an *in silico* approach we demonstrated that tight co-expression of *CACNA1C* and *KCNH2*, against a background of variability of all other ion channels, helps to control the duration of repolarization (**Figure 4**). More importantly, the co-expression of *CACNA1C* and *KCNH2* helps to protect the heart from early after depolarizations when they are exposed to drugs that block *I*_Kr_ (**Figure 5**). Our simulations also suggest that inter-individual differences in pro-arrhythmic responses to *I*_Kr_ drug block can be explained by inter-individual differences in levels of *CACNA1C* expression (**Figure 6**). Finally, our *in vitro* experiments on cardiac myocytes derived from hiPSC lines confirmed that the extent of repolarization prolongation seen with block of *I*_Kr_ was inversely correlated with the level of *KCNH2* mRNA expression (**Figure 8**) and the tight correlation between *I*_Kr_ block-induced FPD prolongation and *I*_CaL_ block-induced repolarization shortening validates the tight coupling between *I*_Kr_ and *I*_CaL_ expression despite marked differences in the absolute levels of expression between different cell lines.

Over the last few years, numerous groups have shown that there is considerable heterogeneity of ion channel expression amongst excitable cells, including neurons (33) and cardiac myocytes (7,19). Furthermore, since the pioneering work of Eva Marder and colleagues, the presence of modules of co-expressed ion channel genes has also been well appreciated (15). These previous studies, however, relied on patch clamp analysis of isolated cells (15,34) and qPCR analysis (15) or single molecule fluorescence *in situ* hybridization (35) of individual ion channel mRNAs, which are highly laborious techniques and so have been restricted to only a few important ion channel genes. The advent of high throughput transcriptomic analyses has greatly facilitated the identification of conserved networks of co-expressed gene modules amongst the entire set of expressed genes (21). By applying meta-analytic approaches to a large number of independent datasets one can more readily discern genuine co-expression signals from noise, as well as explore smaller gene modules (21). In our analysis of a subset of rhythmonome genes within a large library of public RNA Seq datasets, we identified two clusters of genes: a first subset that predominantly affects calcium handling and a second subset that predominantly affects membrane potential (see **Figure 1**). The nodes within each of these clusters that are most closely connected within the rhythmonome relative to all other genes are *KCNH2* and *CACNA1C* (see **Figure 1 figure supplement 1**). This is analogous to a network of networks (36), where the *KCNH2*-*CACNA1C* link provides an interconnection of the two networks. Independent evidence to corroborate an important link between calcium handling and regulation of cardiac action potential duration comes from the large genome-wide association studies (GWAS) of QT interval duration which identified SNPs in a number of calcium handling genes as well as *KCNH2* as being important determinants of QT interval in the population (37).

Due to the large number of associations we were testing for in our network analyses, it is possible that some would occur by chance. That we were able to confirm the presence of at least some of the co-expression modules in an independent dataset, i.e. the hiPSC-CM (see **Figure 2**) provides important corroborative evidence that these co-expression patterns are real and therefore likely to have physiological relevance. It should also be noted that many ion channels important for function in adult cardiac myocytes are not expressed at significant levels in immature cardiac myocytes, such as those derived from hiPSC (e.g. *SCN5A*, *KCNQ1*, *KCNJ12*, *KCND3*, (26)). It is therefore possible that we have underestimated the number of genes within the modules of co-expressed genes in adult cardiac myocytes.

The patterns of co-expressed genes we observed show some important similarities, as well as differences to previous studies. Banyasz et al. (34) and Rees et al. (18) have both demonstrated that the expression of *I*_CaL_ is correlated with the sum of the major repolarizing ion currents in guinea-pig and mouse respectively. However, the molecular players involved in cardiac electrical activity in rodents are quite distinct to humans (38). In humans, the most important determinant of repolarization duration at baseline is *I*_Kr_ (39), whereas in guinea-pig *I*_Ks_ and *I*_Kr_ play equally important roles (34) and in mice the fast component of the transient outward current (*I*_to,f_) and the ultra-rapid delayed rectifier (*I*_Kur_) are the major repolarization currents (18). Thus, there is a common factor between our study showing co-expression of *KCNH2* and *CACNA1C* in human and the previous rodent studies, i.e., all studies show a correlation between *I*_CaL_ and the major repolarizing currents present in that species. Our study, however, is the first to demonstrate the importance of co-expressed genes in human heart tissue. In addition to co-regulation at the level of gene expression, there is also evidence for co-ordinated regulation of modules of ion channels occurring at post-transcriptional levels (35,40). That we still observe an inverse correlation between *KCNH2* mRNA levels and repolarization duration in hiPSC-derived cardiac myocytes, as predicted from our *in silico* analyses, demonstrates that co-regulation at the level of gene expression is still an important determinant of overall cardiac electrical function.

Identifying modules of co-expressed genes are the first step in seeking to understand the logic of complex systems (6). Understanding how such modules impact function in health and disease is the next challenge. In neuronal cells, Marder and colleagues have shown that modules of co-expressed ion channels play an important role in regulating action potential firing patterns (41). Rees and colleagues, have demonstrated that modules of co-expressed depolarization and repolarization currents can help to ensure normal amplitude calcium transients, a critical determinant of overall heart function (18). We have extended these studies to show that in normal heart cells, co-expression of *KCNH2* (repolarization) and *CACNA1C* (depolarization) channels help to maintain the plateau duration of the action potential, which in turn likely contributes to regulating the duration and amplitude of the calcium transient. More importantly, our studies provide the first insights into how patterns of co-expressed ion channel genes influence the hearts response to pathological stimuli.

Sudden death due to abnormalities of cardiac electrical signaling is a major cause of mortality (1). Predicting in advance who is more or less susceptible to sudden cardiac death and therefore warrants prophylactic treatment remains challenging (2). A key to being able to predict who is at greatest risk is understanding why different people respond differently to the same pro-arrhythmic stimulus. Based on the results of our *in silico* studies, validated by *in vitro* analyses of hiPSC-derived cardiac myocytes, we have provided two important insights into the nature of interindividual risk for developing arrhythmias in response to drugs that block *I*_Kr_, the major cause of drug-induced cardiac arrhythmias (14). First, cells with low *G*_CaL_ (and hence low *G*_Kr_ at baseline) exhibited the greatest prolongation of AP duration when exposed to *I*_Kr_ drug block. Second, cells with high *G*_CaL_ (and hence high *G*_Kr_ at baseline) showed greater propensity for development of EADs at moderate levels of *I*_Kr_ drug-block (**Figure 6**). An important implication of the observation that a high *G*_CaL_ increases the susceptibility to EADs in response to drug block of *I*_Kr_ is that the co-administration of an *I*_CaL_ blocker should reduce the risk of EADs (as shown in **Figure 7**). This is consistent with the observation that the administration of magnesium, which is a mild calcium channel blocker (31), is helpful in the acute management of patients with drug-induced *torsades de pointes* (14), and conversely that hypomagnesaemia, which would stimulate *I*_CaL_, can exacerbate *torsades de pointes* (42). It is also consistent with the observation that drugs that block *I*_CaL_ and *I*_Kr_ (e.g., verapamil) are not associated with drug-induced arrhythmias (43) and that verapamil prevented the development of torsades de pointes in rabbit hearts exposed to an *I*_Kr_ blocker (44). However, given that calcium channel blockers are contra-indicated in some ventricular arrhythmias (45), and the likelihood that patients who have drug-induced LQTS may have other underlying cardiac conditions (14), one should be cautious about prescribing calcium channel blockers. Conversely, it would be reasonable to consider using calcium channel blockers to treat patients with LQTS type 2 (i.e. patients with an isolated loss of *I*_Kr_ function) who continue to have cardiac events despite treatment with ß-blockers (46).

In summary, we have demonstrated that meta-analysis of large-scale gene expression data sets is a powerful technique for discerning underlying patterns in gene expression, and that this can provide insights into disease causation at an individual level. Specifically, we have demonstrated that the co-expression of *KCNH2* (*I*_Kr_) and *CACNA1C* (*I*_CaL_) plays an important role in regulating cardiac repolarization both in health and in disease.

## Methods

### Analysis of public RNASeq datasets

An aggregate co-expression gene network was built from public data, similar to that described previously (47). Briefly, 75 human RNA-seq expression experiments (listed in Figure 1 source data 1) that passed quality control and had a least 10 samples (3653 samples in total) were downloaded from the Gemma database (48). Approximately thirty thousand genes were used for the network, limited only to those with Entrez gene identifiers. A co-expression network was generated for each experiment by calculating Spearman’s correlation coefficients between every gene pair and then ranking these values (47). An aggregate gene co-expression network was then generated by averaging across all the individual networks, and re-ranking the final network. This final aggregate network was then used to determine the co-expression ranking between genes that encode for the set of ion channels and calcium handling proteins that determine the shape and duration of the human ventricular AP, the so-called rhythmonome gene subset (see Figure 1 source data 1). Network connectivity of the gene set was measured by comparing the weighted local node degree to the global node degree (49). Node degrees are the sum of the total connections a node (here gene) has within a network. Local node degree refers to the sum of connections (here the ranked correlation) within the rhythmonome gene set, while global node degree is the sum of connections to that gene across all the genes in the network. Code for the analysis is available at https://github.com/sarbal/hERG-cal

### Human induced pluripotent stem cell derived cardiac myocytes

Human iPSC lines, generated from healthy patients by Stanford Cardiovascular Institute Biobank, as previously described (50), were a generous donation from Joseph Wu (Standford Cardiovascular Institute). HiPSC colonies were maintained on Matrigel® (Corning) coated plates in chemically defined medium (mTeSR1™, StemCell technologies), and passaged using Dispase (StemCell technologies). For differentiation, hiPSCs were dissociated by incubating at 37°C for 7 minutes with TrypLE™ Express (ThermoFisher) and seeded at 125000 cells/cm^2^ on a Matrigel® coated 12 well plate, in mTeSR™1 medium supplemented with StemMACS™ Y27632 (Miltenyi Biotec). Once the cells reached greater than 95% confluency, differentiation was initiated using STEMdiff™ Cardiomyocyte (CM) differentiation and Maintenance Kit (StemCell technologies). At day 15, CMs were dissociated by incubation in Collagenase Type I (ThermoFisher) for 45 minutes at 37°C to break up the matrix and then incubated in 0.25% Trypsin with EDTA for 7 minutes at room temperature followed by filtering through a 40 µm cell strainer (51). The CMs were seeded on a Matrigel coated 96-well plate (Greiner Bio-One) and maintained in CM maintenance medium for 10-15 days before they were harvested for mRNA expression analysis. Total RNA was obtained from 40,000 hiPSC-CMs lysed using QIAzol Lysis reagent (Qiagen). The RNA was purified using miRNeasy® Mini Kit (Qiagen), and all samples had RIN values >7.5, and were analysed using Agilent Bioanalyzer pico-chip. RNA samples were hybridised with probes designed to detect 35 known rhythmonome genes using nCounter (NanoString Technologies, see Figure 2 source data 1), which was performed at the Ramaciotti Centre for Genomics (UNSW). For quantitative comparisons of different mRNA species across different runs we normalised mRNA levels, relative to GAPDH, to account for any variation in mRNA loading levels. Raw data for mRNA expression levels are provided Figure 2 -Source Data 1 and Figure 8 -Source Data 2

Extracellular field potentials were recorded from hiPSC-CM monolayers using a Maestro-APEX multi-electrode array (MEA) system (Axion Biosystems, Atlanta, GA, USA) as previously described (52). Briefly, cell monolayers were maintained for 20-30 days in STEMDiff cardiomyocte maintenance media (STEM CELL Technologies), on EStim classic 48 well MEA plates (Axion Biosystems, Atlanta, USA), prior to recording. Monolayers, maintained at 37°C, were paced at 1 Hz and signals digitized at 12.5 kHz using AxIS v2.5.1.10 software (Axion Biosystems, Atlanta, GA, USA). Field potential durations were measured from the peak depolarization spike to the peak of the repolarization wave. After 30 minutes of pacing equilibration, Electrograms were recorded for 5 minutes in control conditions, 5 minutes in the presence of cisapride (200nM), and for 5 minutes in the presence of cisparide + nifedipine (50 nM) and measurements averaged over the last 10 beats of each recording period. This allowed for the possibility that if there was any beat to beat variation this could be averaged out over the ten beats. Twelve wells were plated for each line and all wells from which signals were recorded (n=7−12) were included in the analysis. After the MEA recordings were completed cells were harvested for mRNA analysis, as described above. Raw data for MEA recordings are provided in Figure 8 - Source Data 1.

### Computer modelling

Human cardiac APs were simulated using the endocardial configuration of the O’Hara-Rudy (ORD11) model (27) with key conductances modified as described by Krogh-Madsen et al. (28). The original ORD11 code was adapted to run in the Brain Dynamics Toolbox for Matlab (53). To incorporate population variability in ion channel expression levels the maximum conductance for each current was multiplied by conductance scalar (*G*_x_), that was drawn from a random log-normal distribution (39), with unit mean and variance that was systematically manipulated from 0.05 to 0.5. All models were paced at 1Hz with a stimulus of −40 mV and duration of 1 ms and allowed to equilibrate for 300 beats. We then analysed the next four beats (to allow for the possibility of development of alternans) after the equilibration stage. The peaks in those APs were identified using the Matlab *findpeaks* function. Individual beats were classified as ectopic if they had secondary peaks that were separated by more than 100 ms. The set of four beats were further classified as *alternans* if the profile of any of the APs deviated from each other by more than 1 mV at any time point. In a second set of simulations, we repeated the same method as above except that the random multipliers applied to both *I*_CaL_ and *I*_Kr_ were identical. This case we denote co-expression whereas the former case we denote independent expression. Code for the AP modeling is available at https://git.victorchang.edu.au/projects/CC/repos/ordcoreg.

## Supporting information

Figure 8 source data 1 summary

## Acknowledgements

This work was supported by the Victor Chang Cardiac Research Institute Innovation Centre, funded by the NSW Government. We also thank Terry Campbell, Dan Roden and Raj Subbiah for helpful discussions.

## Supplementary Files

**Figure 3 supplementary Movie 1**: Simulations of 1000 APs with all conductance scalars allowed to vary independently

**Figure 4 supplementary Movie 1**: Simulations of 1000 APs with *G*_Kr_ and *G*_CaL_ scalars coupled but all other conductance scalars allowed to vary independently

## Data Sets

Figure 1 – Source Data 1.xlsl

Sheet 1: List of GEO datasets analysed
Sheet 2: Co-expression rankings for rhythmonome genes
Sheet 3: modules of co-expressed genes
Figure 2 – Source Data 1: nanostring datasets: GEO: TBA

Sheet 1: Metadata
Sheet 2: normalized mRNA read counts
Raw data files (uploaded to GEO)
Figure 8 – Source Data 1: Multi-electrode array datasets

1a: MEA guide.txt (description of contents of raw data files and how to extract the data).
1b: plate1_baseline.mat
1c: plate1_cis.mat
1d: plate1_nif.mat
1e: plate2_baseline.mat
1f: plate2_cis.mat
1g: plate2_nif.mat
Figure 8 – Source Data 2: nanostring datasets: GEO: TBA

Sheet 1: Metadata
Sheet 2: normalized mRNA read counts
Raw data files (uploaded to GEO)

## Software Code

Figure 1,2; meta data analysis scripts: https://github.com/sarbal/hERG-cal

Figures 3–7: Baseline code and BD Toolbox software package used to generate figures 3–7: https://git.victorchang.edu.au/projects/CC/repos/ordcoreg.

---

**Figure 1 - figure supplement 1:**
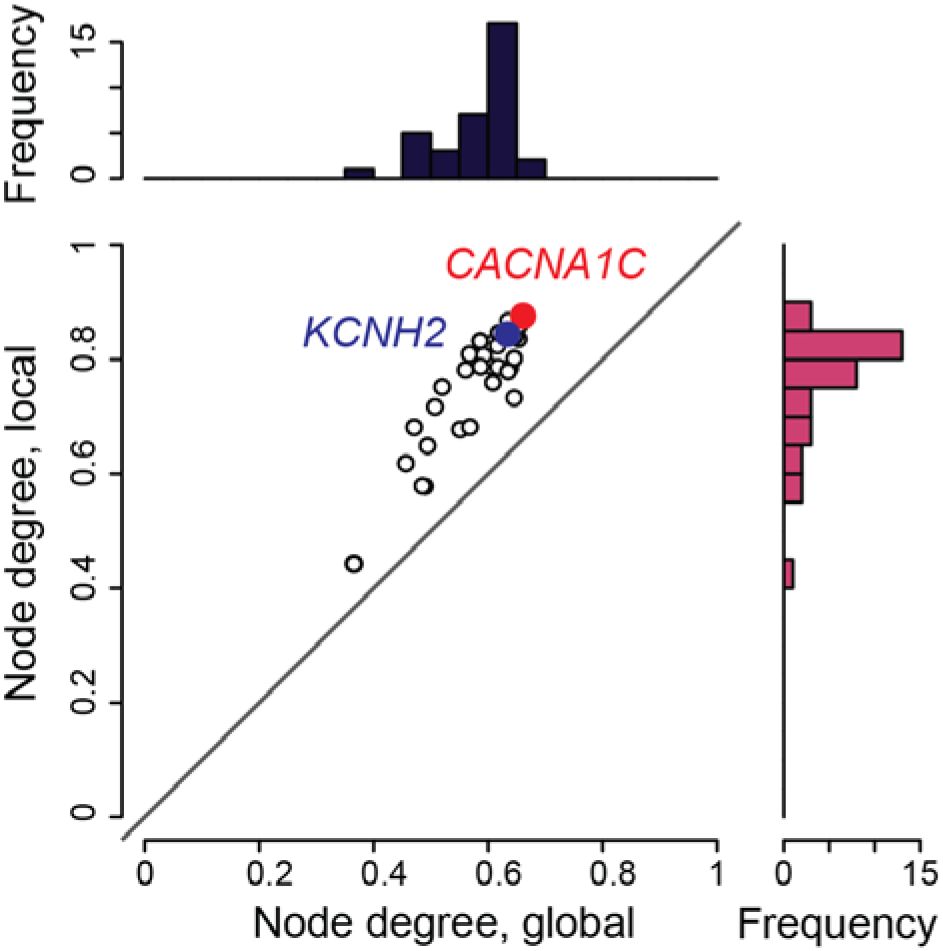
Network connectivity for each rhythmonome gene. Network connectivity for each rhythmonome gene within the local rhythmonome network (y-axis) versus its connectivity within the whole genome (x-axis). All the rhythmonome genes are more highly connected within the local network (i.e., all the points lie above the line of identity, p~5.7e^−14^). **CACNA1C** is highlighted in red and **KCNH2** in blue.

**Figure 3 Movie supplement 1:**
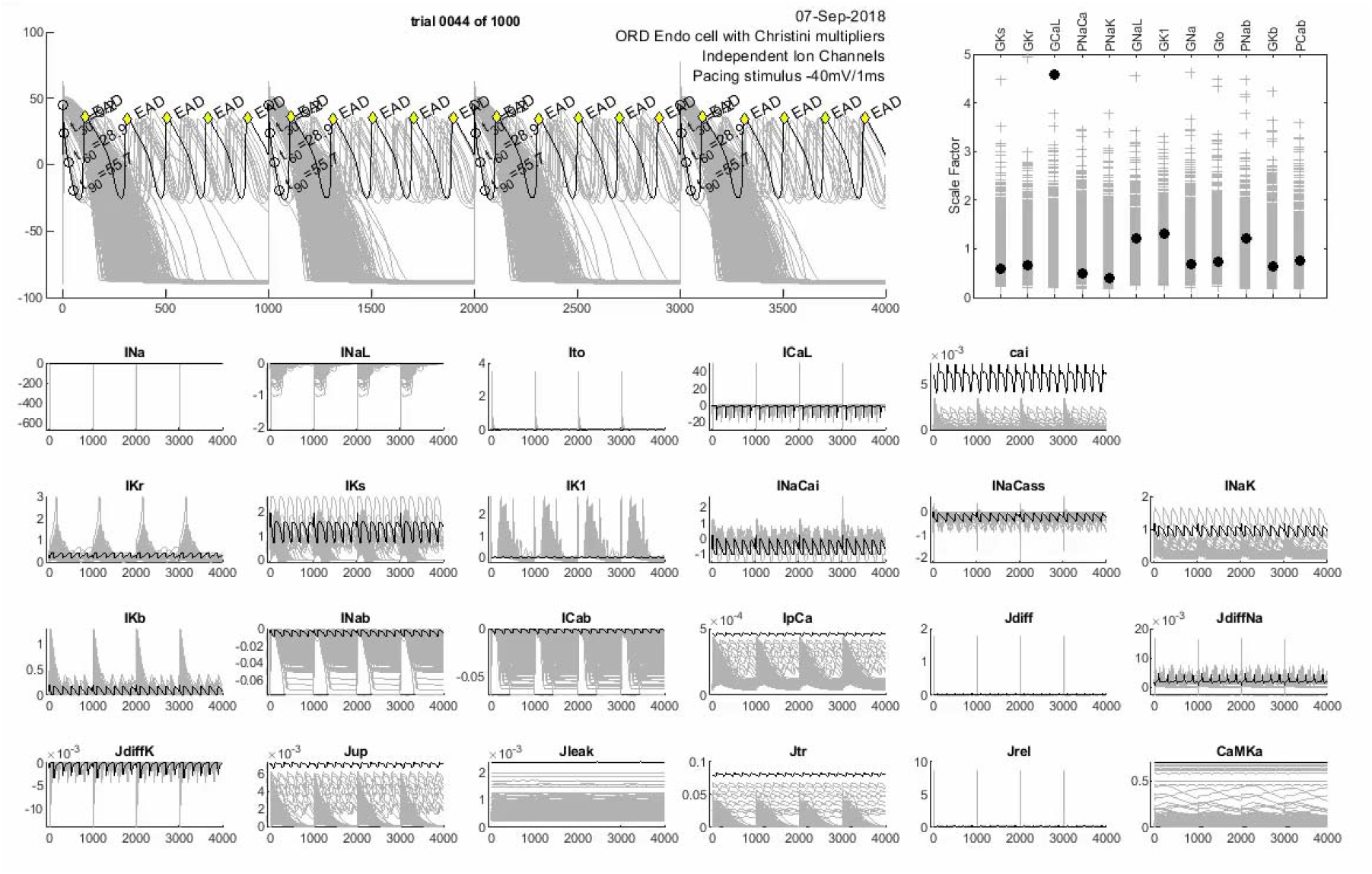
Last 4 beats for the population of 1000 simulated APs.

**Figure 4 figure supplement 1:**
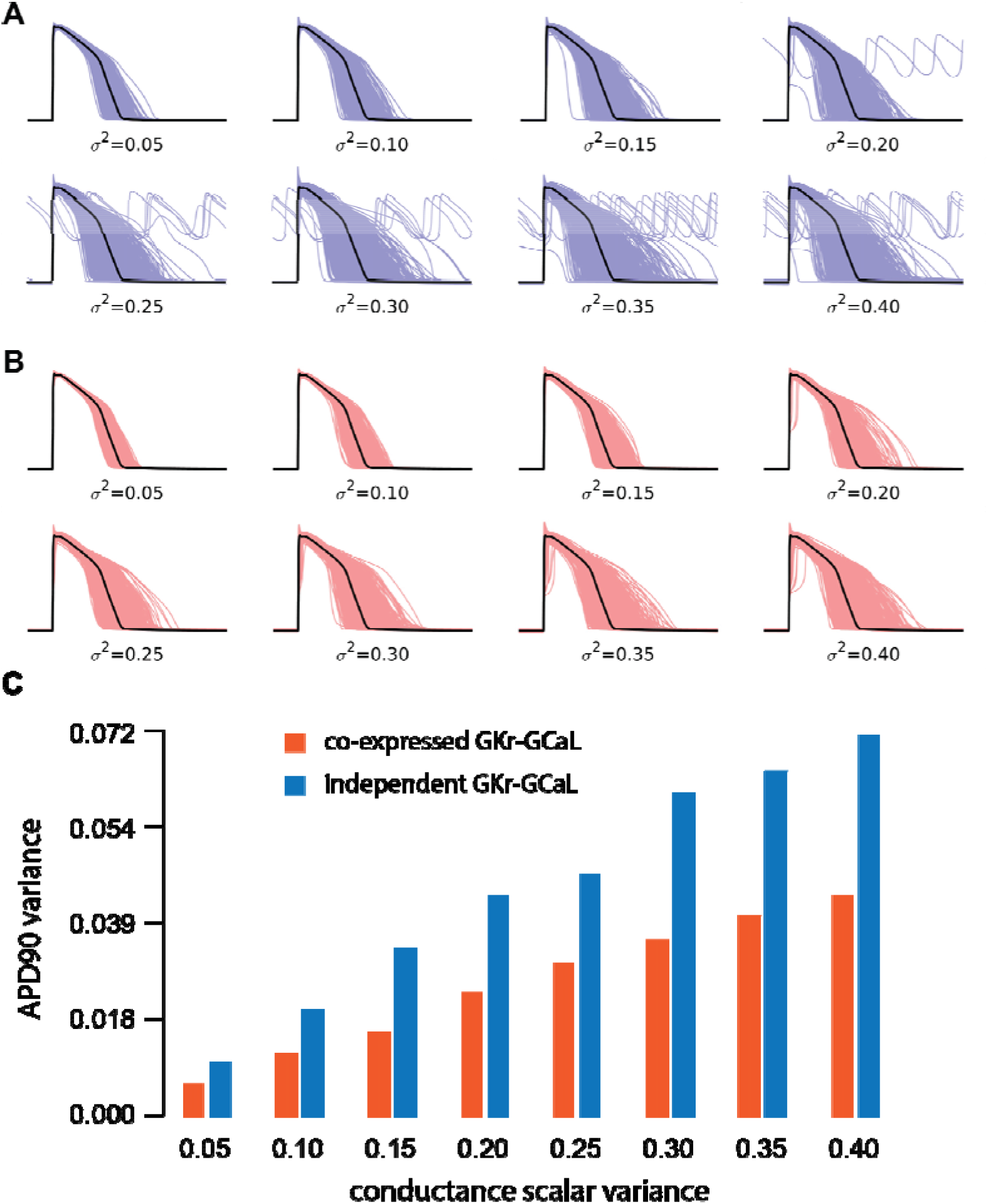
Population models of ventricular APs with different levels of scalar variance. **A-B**. Population of models with variance systematically increased from σ2 = 0.05 to 0.40 for independent conductance scalars for *G*_Kr_ and *G*_CaL_ **(A)**, or co-expression conductance scalars for *G*_Kr_ and *G*_CaL_ **(B)**. As the variance increases, the number of cells that develop EADs increases in the independent series. Conversely, for the co-expression models, there are no EADs even at variances up to 0.4. **C.** Variance of the APD_90_ distributions for each set of conductance scalar inputs for independent (blue) and co-expression (red) of *I*_Kr_ and *I*_CaL_. Note that the variance of the APD_90_ distribution was always greater for the independent simulations (p<0.008 between groups, paired t-test)

**Figure 4 Movie supplement 1.**
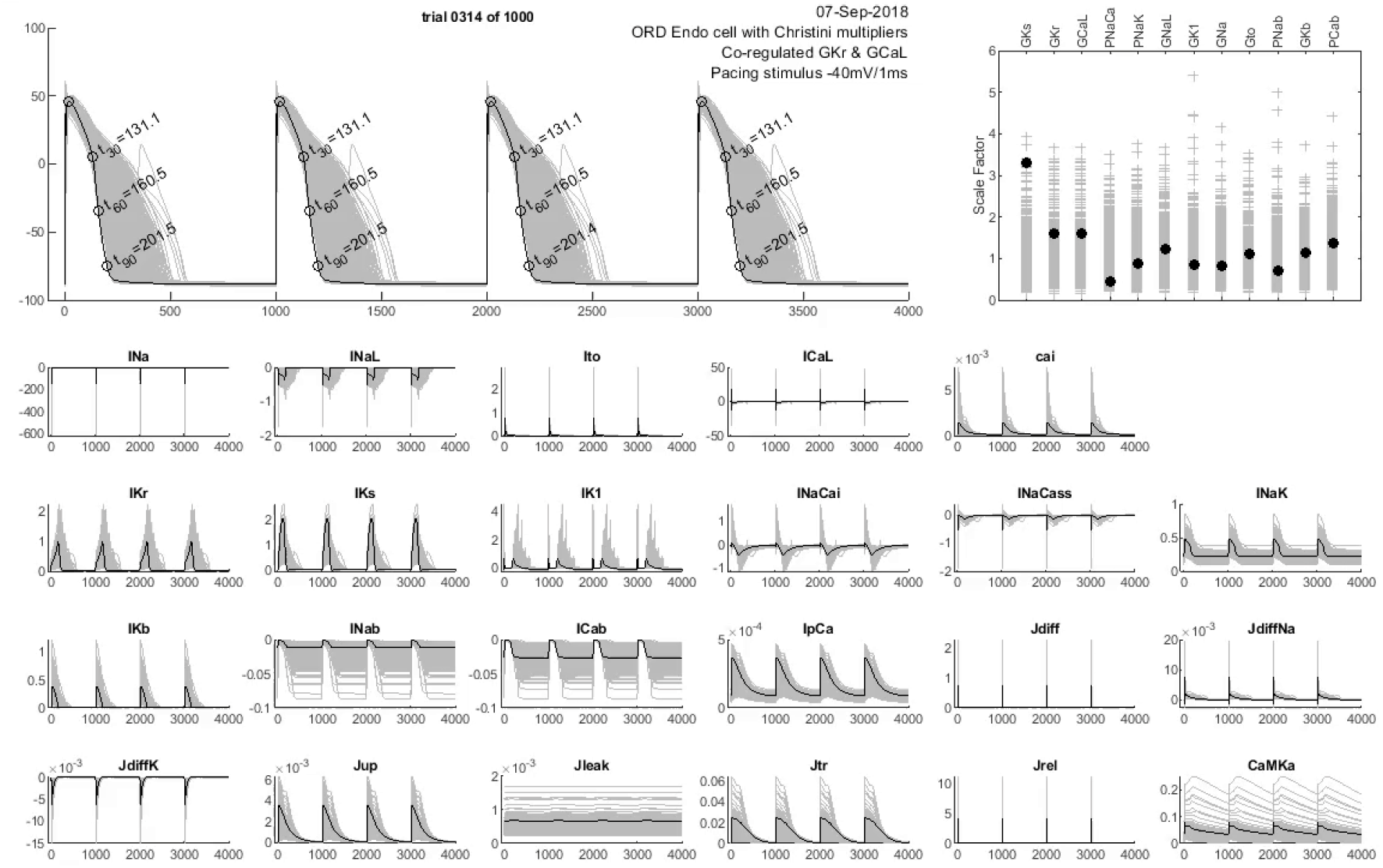

**Figure 6: figure supplement 1:**
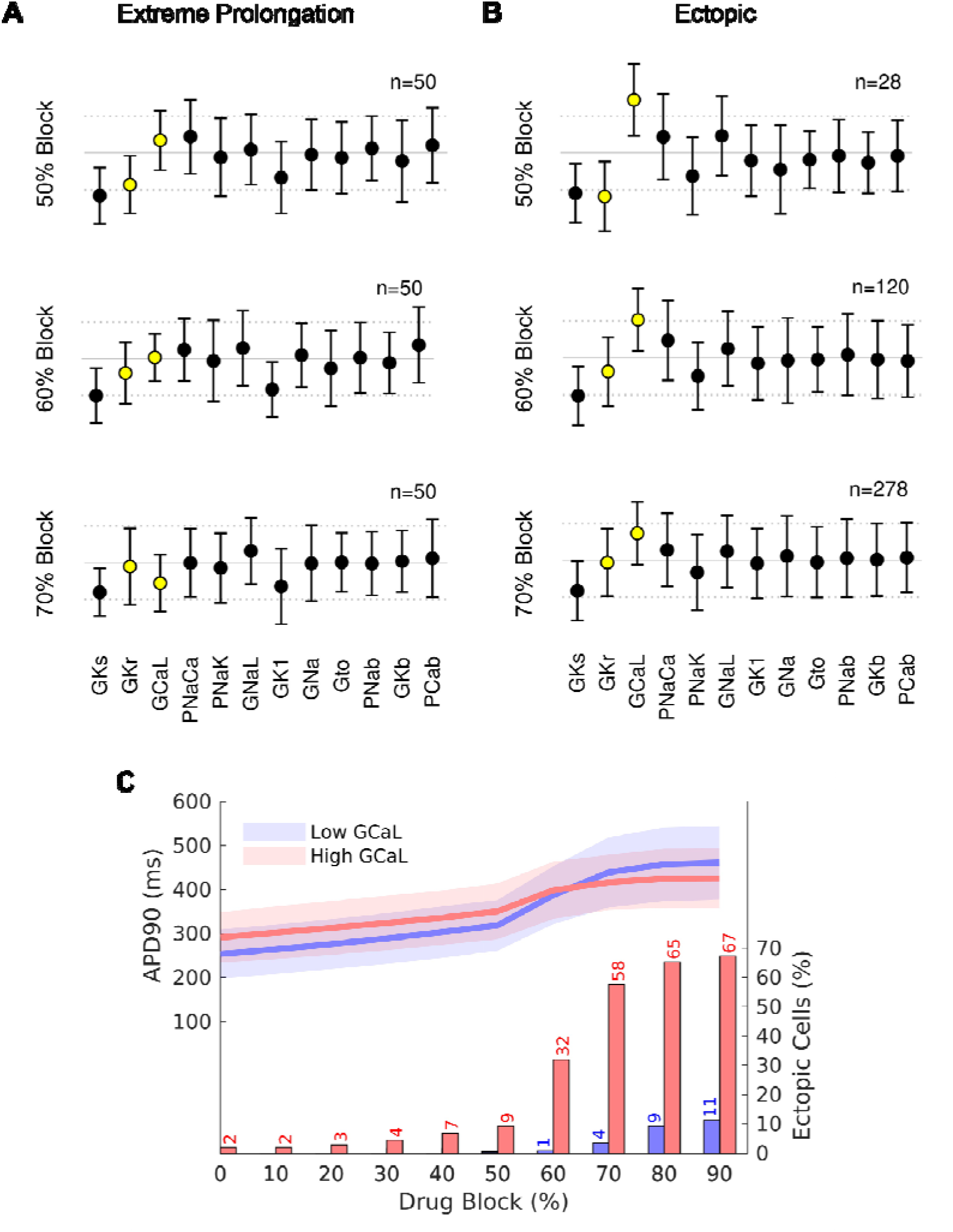
High *G*_CaL_ increases risk of EAD formation at moderate *I*_Kr_ block even when *G*_CaL_ and *G*_Kr_ are varied independently. Conductance scalars (mean ± SD) that give **A.** The 50 most extreme APD values or **B.** EADs for the population of cells with independent regulation of *G*_Kr_ and *G*_CaL_. **C.** Plot of APD_90_ values for the highest quartile of *G*_CaL_ (red) and lowest quartile of *G*_CaL_ (blue) versus % drug block of *I*_Kr_. The continuous line shows mean and shaded regions show ± 1 standard deviation. The column graph shows the % of cells that develop EADs at each level of drug block.

