## Supplementary material for "Co-expression of calcium channels and delayed rectifier potassium channels protects the heart from proarrhythmic events": Figure 8 source data 1 summary

Multielectrode array data related to Figure 8 is captured in the files:
plate1_baseline
plate1_cis
plate1_nif
plate2_baseline
plate2_cis
plate2_nif
where the suffix 'baseline', 'cis' and 'nif' correspond to data for absence of drug, 10 mins after addition of 200 micmolar cisapride, and 10 mins after addition of 50 micmolar nifedipine respectively.

plate1 files contain data from cell lines 142C1, 273C1 and 274C1.
plate2 files contain data from cell lines 8C1, 15C1 and 479C1.


Each file contains a structure 'FileData' that contains the filename, experimental notes (Experiment ID, Investigator, and Description), the file version number of the file, the plate map, and any tag events associated with the file.

Where recorded, experimental information is stored in strings, which can be accessed from the loaded file by calling:

Investigator = FileData.Notes.Investigator;
ExperimentID = FileData.Notes.ExperimentID;
Description = FileData.Notes.Description;
PlateMap = FileData.PlateMap;
HardwareSettings =  FileData.DataSets.HeaderExtension.Description


Each file also contains 3 cell arrays: AllWells_AvgBeats, AllWells_UnaveragedTime, and AllWells_UnaveragedData

AllWells_UnaveragedData is a 6 X 8 cell array corresponding to the number of (Well Rows) x (Well Columns). Each cell contains raw data from each of the 16 electrodes in the well for the ten most stable beats (determined by beat rate variability) that were analsysed.
AllWells_UnaveragedTime is a 6 X 8 cell array corresponding to the number of (Well Rows) x (Well Columns). Each cell contains the timeseries corresponding to data in AllWells_UnaveragedData
AllWells_AvgBeats is a 6 X 8 cell array corresponding to the number of (Well Rows) x (Well Columns). Each cell contains the averaged waveform for the ten beats included in AllWells_UnaveragedData, from which FPD measurements were made.
